# Whole-brain modeling of dynamic causal circuits in human cognition using amortized variational inference

**DOI:** 10.64898/2026.08.03.742253

**Authors:** Byeongwook Lee, Louis Rouillard, Leticia Levin Diniz, Linjing Jiang, Luca Ambrogioni, Srikanth Ryali, Nicholas Branigan, Percy Mistry, Weidong Cai, Demian Wassermann, Vinod Menon

## Abstract

Understanding dynamic mechanisms underlying cognition remains a major challenge in human neuroscience. Here, we develop, validate, and apply *Multivariate Dynamical Systems Identification with Amortized Variational Inference* (MDSI-AVI), a novel computational framework designed to address critical challenges in capturing asymmetric, context-dependent, whole-brain directed interactions while accounting for regional hemodynamic response variability in fMRI data. MDSI-AVI leverages simulation-based inference through forward and reverse variational inference to address the limitations of conventional variational methods in high-dimensional settings. By averaging over uncertainty in hemodynamic response parameters using forward simulation, MDSI-AVI provides well-calibrated posteriors of directed connectivity that scale efficiently to networks with hundreds of nodes. Applied to Human Connectome Project data (*N*=728), MDSI-AVI reveals new insights into working memory mechanisms, identifying the dorsal anterior insula as a critical hub influencing activity at the whole-brain level. We demonstrate task-dependent modulation of causal influences, where the salience network drives frontoparietal network activity, which differentially influences the default mode and sensorimotor networks depending on working memory load. These whole-brain causal interactions distinguish task conditions with high accuracy and predict working memory performance. Our framework demonstrates reproducible results across whole-brain parcellations, establishing MDSI-AVI as a robust tool for advancing our understanding of circuit dynamics in cognition and disease.

## INTRODUCTION

Understanding whole-brain dynamics is fundamental to human cognition ^1,2^, yet existing analytical methods cannot fully capture the complex causal interactions spanning the entire brain. A critical challenge in human neuroscience is characterizing how brain-wide networks dynamically interact during cognitive tasks, and particularly how these interactions vary with cognitive demands. Here, we develop Multivariate Dynamical Systems Identification with Amortized Variational Inference (MDSI-AVI), a novel computational framework designed to address critical challenges in capturing asymmetric, context-dependent, whole-brain directed interactions while accounting for regional hemodynamic response variability in fMRI data.

Analyzing causal control circuits at the whole-brain scale promises to elucidate how asymmetries in directed influence enable specific brain regions or networks to exert control over others ^2^. This knowledge is crucial for neuromodulation studies, which are increasingly used to treat both psychiatric and neurological disorders ^3–5^.

A quintessential example of a cognitive function that operates at the whole-brain level is working memory, the ability to temporarily hold and manipulate information in the mind ^6–10^. Working memory is essential for most cognitive processes, and its impairments are prominent in many psychiatric and neurological disorders including schizophrenia, autism, major depression, and epilepsy ^11–14^. Despite significant advances in identifying brain regions and exploring interactions among specific neural circuits ^15–17^, our understanding of the dynamic mechanisms operating at the whole-brain level remains limited.

Traditional studies of working memory and many other higher-order cognitive processes have often concentrated on a handful of brain regions such as the dorsolateral prefrontal cortex ^18–21^ and posterior parietal cortex ^22,23^. However, these approaches cannot capture the emergent properties that arise from whole-brain interactions. This knowledge gap is particularly evident when considering how the entire brain orchestrates working memory operations. It has been suggested that working memory operates primarily through dynamic interactions at the circuit level, rather than sustained activation in individual regions ^24^. Understanding the neural basis of working memory increasingly involves examining dynamic functional connectivity across the entire brain ^15–17,24^. Despite significant advances, our understanding of whole-brain circuit mechanisms remains limited, as prior studies have primarily focused on interactions among a small number of pre-determined brain regions due to methodological constraints.

A key challenge has been that previous studies have lacked the computational capability to comprehensively model causal multivariate dynamics across the entire brain ^2^. This selective approach often introduces biases by predominantly favoring well-researched brain areas. These biases impede a comprehensive understanding of brain circuitry by restricting the exploration of less-studied regions and potentially overlooking novel or unexpected neural connections.

Moreover, in multivariate models, the influence of brain areas excluded from the analysis is often neglected ^7,25,26^. This oversight can lead to incomplete or skewed interpretations of neural interactions, as the effects of these omitted regions on overall brain network dynamics remain unexplored.

The comprehensive coverage afforded by fMRI provides a unique vantage point for discovering context-sensitive direct and indirect asymmetric directed interactions across the entire brain. This whole-brain perspective allows for data-driven and unbiased exploration of potentially unexpected interactions throughout the brain. Unlike invasive technologies such as calcium imaging, two-photon microscopy, or emerging neuropixel technologies, which offer limited coverage, fMRI can non-invasively capture brain activity patterns. Moreover, advances in sub-second temporal resolution with fMRI are pushing its capabilities even further ^27,28^. The accessibility of large datasets from projects such as the Human Connectome Project (HCP) with high temporal sampling provides unique opportunities to examine the stability and robustness of whole-brain findings, addressing concerns about reproducibility in human neuroscience research ^29,30^ which can arise from varied approaches to specifying brain regions of interest.

However, analysis of dynamic interactions at the whole-brain level presents formidable challenges. The primary challenge is scaling up the analysis to encompass the entire brain due to the high-dimensional parameter space involved. Joint multivariate modeling is essential but complex ^31^. Additionally, accounting for variations in the hemodynamic response function (HRF) across the whole brain ^32–35^ is crucial, as these variations can significantly impact the accuracy of estimates ^7,36,37^. This issue becomes more pronounced when modeling the entire brain at high spatial resolution. Furthermore, causal interactions among brain regions evolve dynamically with the experimental context, necessitating sophisticated modeling techniques to capture these whole-brain dynamics accurately (**Figure 1A**).

**Figure 1.**
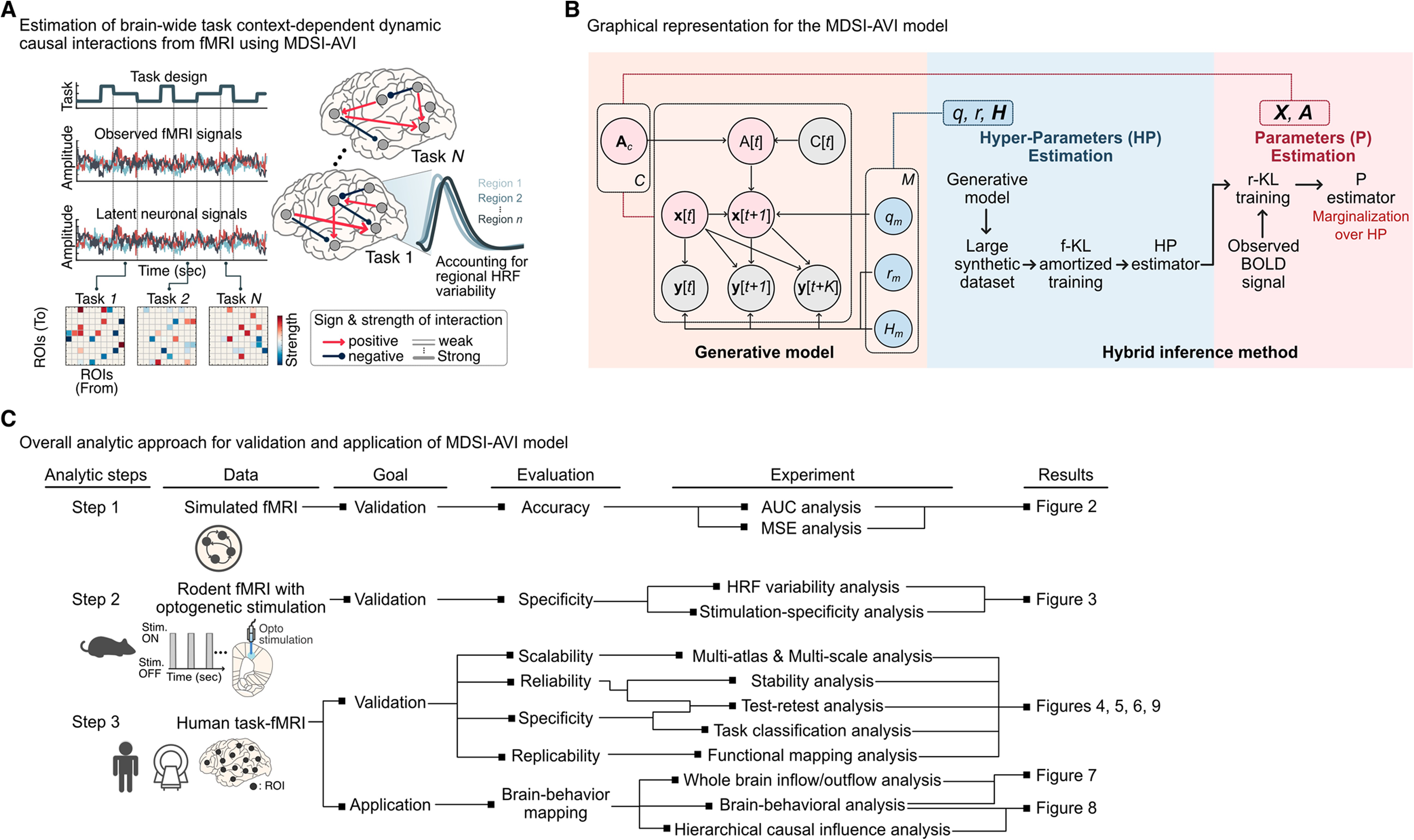
MDSI-AVI framework and validation strategy for whole-brain causal modeling. **(A)** Conceptual overview. MDSI-AVI estimates task-dependent causal interactions across the entire brain from fMRI data while accounting for regional variations in hemodynamic responses. **(B)** Model architecture and inference strategy. MDSI-AVI employs a hybrid two-stage approach to overcome scalability and identifiability challenges in whole-brain causal modeling. Stage 1 (Amortized Inference): An encoder and normalizing flow learn posterior distributions of hyperparameters – including latent noise (**q**), observation noise (**r**), and regional hemodynamic response parameters (**H**) – from observed fMRI signals. This step addresses the fundamental many-to-one mapping problem from neural activity to fMRI observations by capturing uncertainty in these parameters. Stage 2 (Subject-specific Inference): Individual-level inference minimizes reverse Kullback-Leibler divergence to estimate latent neural activity (**X**) and directed connectivity matrices (**A**), conditioned on the learned hyperparameter distributions. This principled uncertainty marginalization significantly improves estimation of individual causal brain dynamics and enhances prediction accuracy. **(C)** Three-stage validation and application strategy. We systematically validated MDSI-AVI using: (1) simulated fMRI data with known ground truth to assess accuracy; (2) optogenetic rodent fMRI data to evaluate specificity for stimulus-evoked causal interactions; and (3) human n-back working memory task data to demonstrate scalability, reliability, and replicability across sessions and brain atlases. Finally, we examined the relation between causal network interactions and working memory performance.

Recent studies have highlighted the importance of large-scale brain networks, particularly the Salience Network (SN), Frontoparietal Network (FPN), and Default Mode Network (DMN), in supporting working memory ^38–42^. These networks form part of a larger whole-brain architecture: the SN, anchored in the anterior insula and dorsal anterior cingulate cortex, is thought to play a crucial role in detecting behaviorally relevant stimuli and initiating brain systems involved in cognitive control ^43,44^; the FPN, including the dorsolateral prefrontal cortex and posterior parietal cortex, coordinates the active maintenance and manipulation of information ^45–49^; and the DMN, typically deactivated during cognitive tasks, has been implicated in internal mentation and may play a role in working memory through its whole-brain interactions as a global signaling hub ^39,41,50,51^. However, the precise dynamics of how these networks interact within the broader context of whole-brain organization to support working memory, particularly under varying cognitive loads, remain poorly understood.

MDSI-AVI employs a state-space modeling framework that separates the inference problem into two key components: a’state equation’ capturing unobservable neural dynamics and an’observation equation’ linking these dynamics to measured fMRI signals ^37,52,53^. To handle the computational challenges of whole-brain analysis, MDSI-AVI introduces a novel hybrid inference approach that combines forward and reverse variational methods (**Figure 1B**). The forward component uses amortized variational inference to estimate region-specific hemodynamic responses and their uncertainties ^54,55^, while the reverse component employs a scalable gradient-based method to estimate directed causal interactions between brain regions. This separation enables MDSI-AVI to effectively account for regional variations in hemodynamic responses while efficiently estimating causal interactions among hundreds of brain regions. The hybrid approach is particularly powerful because it maintains computational tractability while properly accounting for uncertainties in both the hemodynamic responses and the underlying neural interactions (see **Methods and Supplementary Methods** for details).

We demonstrate that MDSI-AVI provides an elegant analytical solution for computing the posterior distributions and deriving sparse, interpretable causal connectivity estimates. We validate MDSI-AVI using extensive neural simulations and optogenetic stimulation experiments designed to test whole-brain modeling capabilities (**Figure 1C**). We then apply our approach to a large HCP dataset (*N*=728), leveraging its unique sample size to investigate dynamic causal interactions during working memory at the whole-brain level (**Figure 1C**). By providing a comprehensive multi-scale characterization of whole-brain dynamic causal interactions during working memory, our study aims to overcome limitations of prior research that primarily focused on pre-selected regions of interest ^7,56,57^ and to advance our understanding of the distributed whole-brain dynamics underlying this essential cognitive function. Our approach not only addresses critical needs in human brain research by pioneering the development and validation of new robust tools for identifying dynamic causal interactions at the whole-brain level, but also paves the way for a deeper understanding of how the entire brain dynamically coordinates signaling to support working memory processes.

## RESULTS

### Validation of MDSI-AVI using small-scale simulated fMRI data

To evaluate the performance of MDSI-AVI in identifying dynamic causal interactions, we first tested it on simulated data with known connectivity. We used simulated fMRI data from a benchmark of nine networks (5-10 nodes) introduced by Sanchez-Romero et al. ^58^, varying in density and number of cycles (**Figure 2A**). For each network configuration, 60 individual datasets were generated with introducing controlled variation in connection magnitudes and node-wise hemodynamic response functions (HRFs) (see **Methods** for details). This setup enabled us to systematically evaluate our model’s ability to recover ground-truth connectivity under HRF variability.

**Figure 2.**
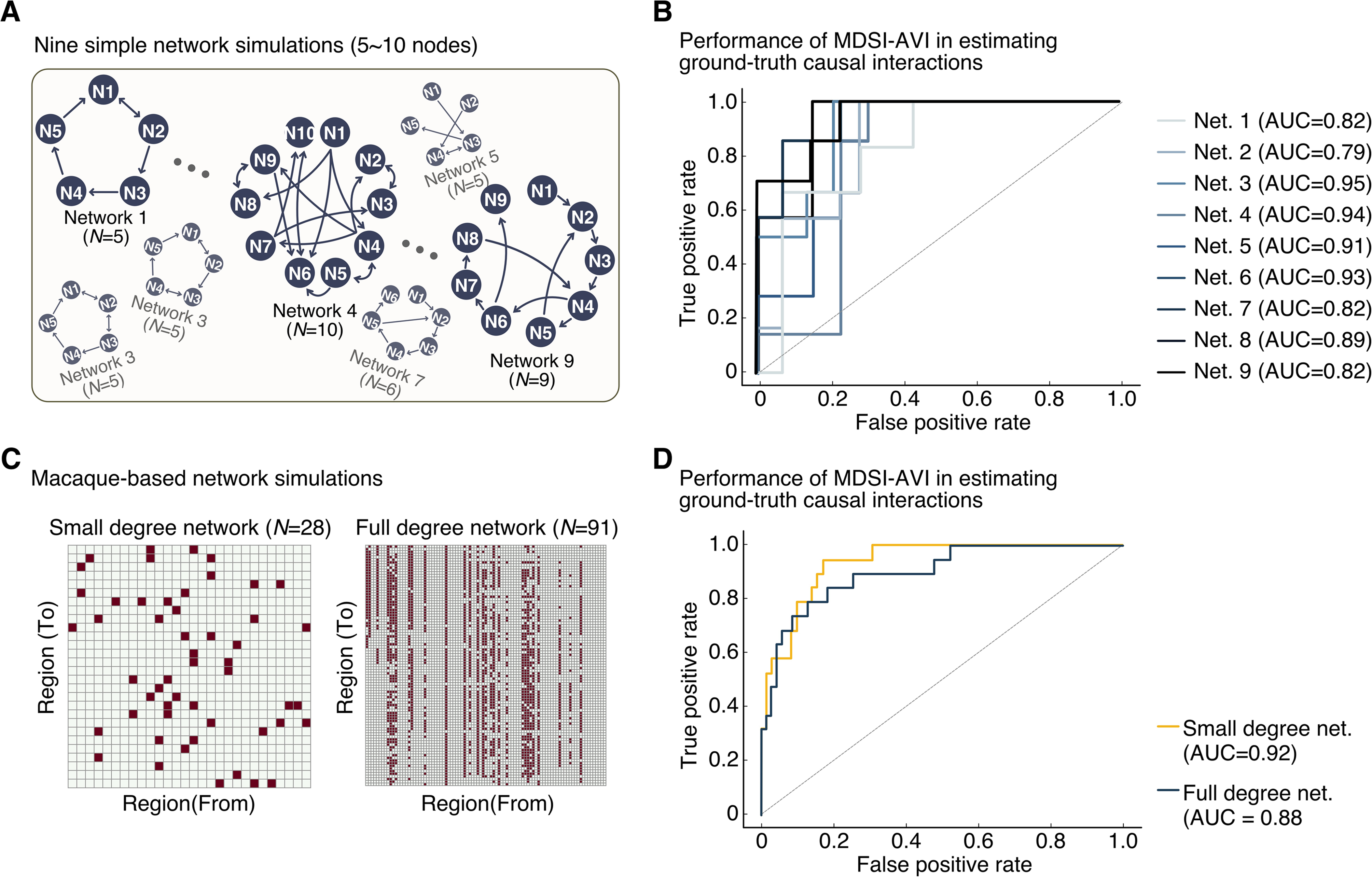
MDSI-AVI demonstrates high accuracy in recovering ground-truth connectivity from simulated fMRI data. **(A)** Simple benchmark networks. Nine synthetic networks varying in density and complexity serve as ground-truth connectivity patterns for initial validation. **(B)** Performance on simple networks. ROC curves demonstrate that MDSI-AVI consistently performs well above chance (gray diagonal) across all network configurations, achieving a mean AUC of 0.87. This establishes reliable edge detection capability across diverse network topologies. **(C)** Macaque connectome-based networks. Two realistic benchmarks derived from the complete macaque anatomical connectome: a smaller network (28 nodes, 52 edges) and a larger, more complex network (91 nodes, 1,615 edges). **(D)** Performance on realistic large-scale networks. MDSI-AVI maintains excellent performance on neurobiologically realistic networks, achieving a mean AUC of 0.91. The sustained high accuracy across both small and large-scale networks demonstrates the method’s scalability and robustness for whole-brain causal modeling applications.

We evaluated the performance of MDSI-AVI using two complementary metrics: (1) the area under the receiver operating characteristic curve (AUC), which assesses the binary classification of connection presence, and (2) the mean absolute percentage error (MAPE), which quantifies deviations between estimated and true connection strengths. MDSI-AVI achieved strong performance across all nine networks, with a mean AUC of 0.87 (**Figure 2B**) and mean MAPE of 13.5% (**Supplementary Figure 1** and **Supplementary Table 1).** These results demonstrate that MDSI-AVI can uncover underlying connectivity with high accuracy in small-scale simulated networks.

### Validation of MDSI-AVI using whole-brain simulated fMRI data and the macaque anatomical connectome

Next, we examined MDSI-AVI’s performance on larger-scale, more realistic simulated benchmarks based on the whole-brain macaque anatomical connectome ^59^, one with 28 nodes and 52 directed edges and another with 91 nodes and 1,615 directed edges (**Figure 2C**). MDSI-AVI accurately uncovered the underlying causal interactions in both networks, obtaining a mean AUC of 0.90 and mean MAPE of 10.5% across the two networks (**Figure 2D** and **Supplementary Table 1**). These results demonstrate the robustness, accuracy, and scalability of MDSI-AVI in identifying underlying causal interactions in neurobiologically realistic whole-brain simulated data.

### Validation of MDSI-AVI using opto-fMRI stimulation

To further validate MDSI-AVI’s ability to uncover dynamic causal interactions, we used whole-brain rodent optogenetic fMRI (opto-fMRI) data acquired during stimulation of the primary motor cortex (M1) ^53^ (**Figure 3A**). Opto-fMRI allows for probing the direct effects of in-vivo brain stimulation to characterize changes in dynamic causal interactions between stimulated and downstream target brain regions. fMRI time series were extracted from M1, the thalamus (a downstream target), and the insula (a non-activated control region) (**Figure 3B**). MDSI-AVI captured regional HRF variability across the three regions of interest in all three rodents (**Figure 3C**) and exclusively and specifically identified an increased causal interaction from M1 to thalamus during ON-stimulation, with no significant increases observed in other connections, compared to OFF-stimulation (**Figure 3D**, *p* < 0.01, Bonferroni correction). This causal drive from M1 to thalamus was detected with the addition of insula as a control region in all three rats. These results demonstrate the sensitivity of MDSI-AVI in identifying dynamic causal interactions and its ability to accurately represent latent neural activity, reflecting true neural dynamics rather than mere blood flow changes.

**Figure 3.**
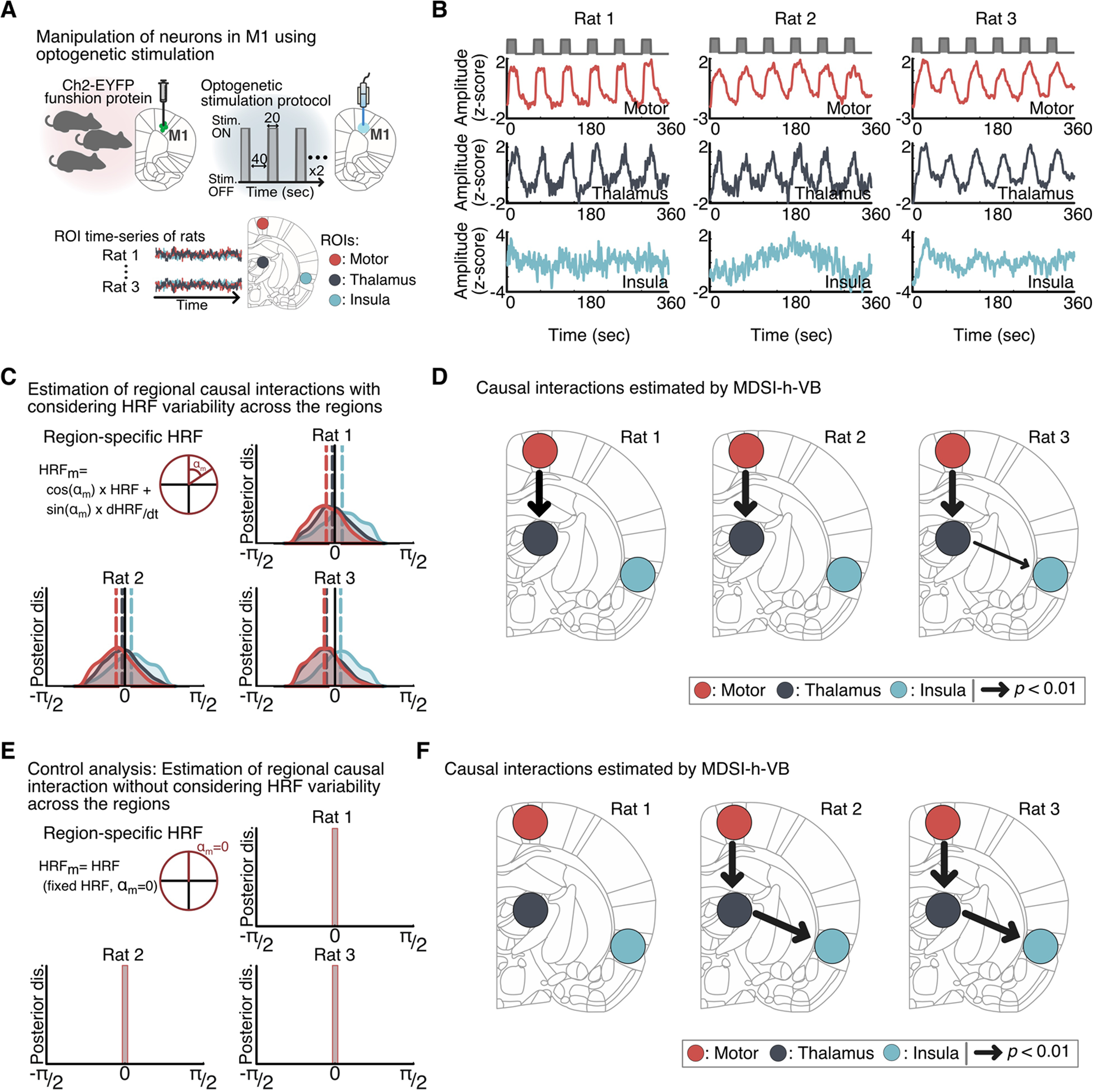
MDSI-AVI accurately detects stimulus-evoked causal interactions in optogenetic fMRI experiments. **(A)** Experimental design. Primary motor cortex (M1) was targeted with adeno-associated virus expressing Channelrhodopsin2-EYFP, with chronically implanted optical fibers positioned for precise stimulation. **(B)** Representative fMRI signal time courses from three experimental animals showing stimulus-evoked responses. **(C-D)** Causal interactions with regional HRF modeling. MDSI-AVI with HRF variability modeling specifically identified the expected M1→thalamus causal connection in rats 1 and 2 (*p* < 0.01, Bonferroni corrected), with rat 3 showing an additional thalamus→insula connection. **(E-F)** Control analysis without HRF modeling. When regional HRF variability was ignored, the method failed to consistently detect the true connections, demonstrating that proper HRF modeling is essential for accurate causal inference.

To evaluate the necessity of accounting for regional HRF variability, we conducted a control analysis using a canonical HRF, instead of region-specific HRFs (**Figure 3E**). We found that the M1 to thalamus connection was not consistently detected across rats, and a spurious thalamus to insula connection was detected in two out of three rats (**Figure 3F**). This result suggests that accounting for regional HRF variability is crucial for reliably estimating correct underlying causal interactions.

### Whole-brain modeling of causal circuit dynamics in human working memory: scalability and reliability of MDSI-AVI

Having demonstrated MDSI-AVI’s ability to uncover underlying connectivity with high accuracy and capture HRF heterogeneity across different brain regions in simulated data as well as in actual brain data, we next examined its application to brain-wide causal circuit dynamics in human working memory. We leveraged DiFuMo atlases at different granularities ^60^ to test the robustness, scalability, and functional interpretability of MDSI-AVI in extracting brain-wide causal circuit dynamics underlying working memory. The DiFuMo atlas provides multiple fine-grained brain-wide parcellations of both cortical and subcortical areas at different levels of granularity. This feature allowed us to test the scalability of MDSI-AVI in estimating whole-brain causal interactions across a large number of brain regions. We leveraged large samples of data (*N*=728) from the Human Connectome Project (HCP) to validate our findings using replicability and stability analyses.

We used the DiFuMo atlas with dimensions of 64, 128, and 256, and computed the average task-fMRI time series across all voxels in each node at each scale (**Figure 4A**). We then estimated whole-brain causal interactions for each dimension. MDSI-AVI identified asymmetric connectivity patterns exhibiting significant dynamic causal interactions under rest, 0-back, and 2-back task conditions (*p*<0.01, FDR-corrected, two-sided paired *t*-test) across each dimension (**Figure 4B**). MDSI-AVI took approximately 250, 300, and 500 seconds, on each participant, to infer directed connectivity patterns in 64, 128, and 256 dimensions, respectively (**Figure 4C**).

**Figure 4.**
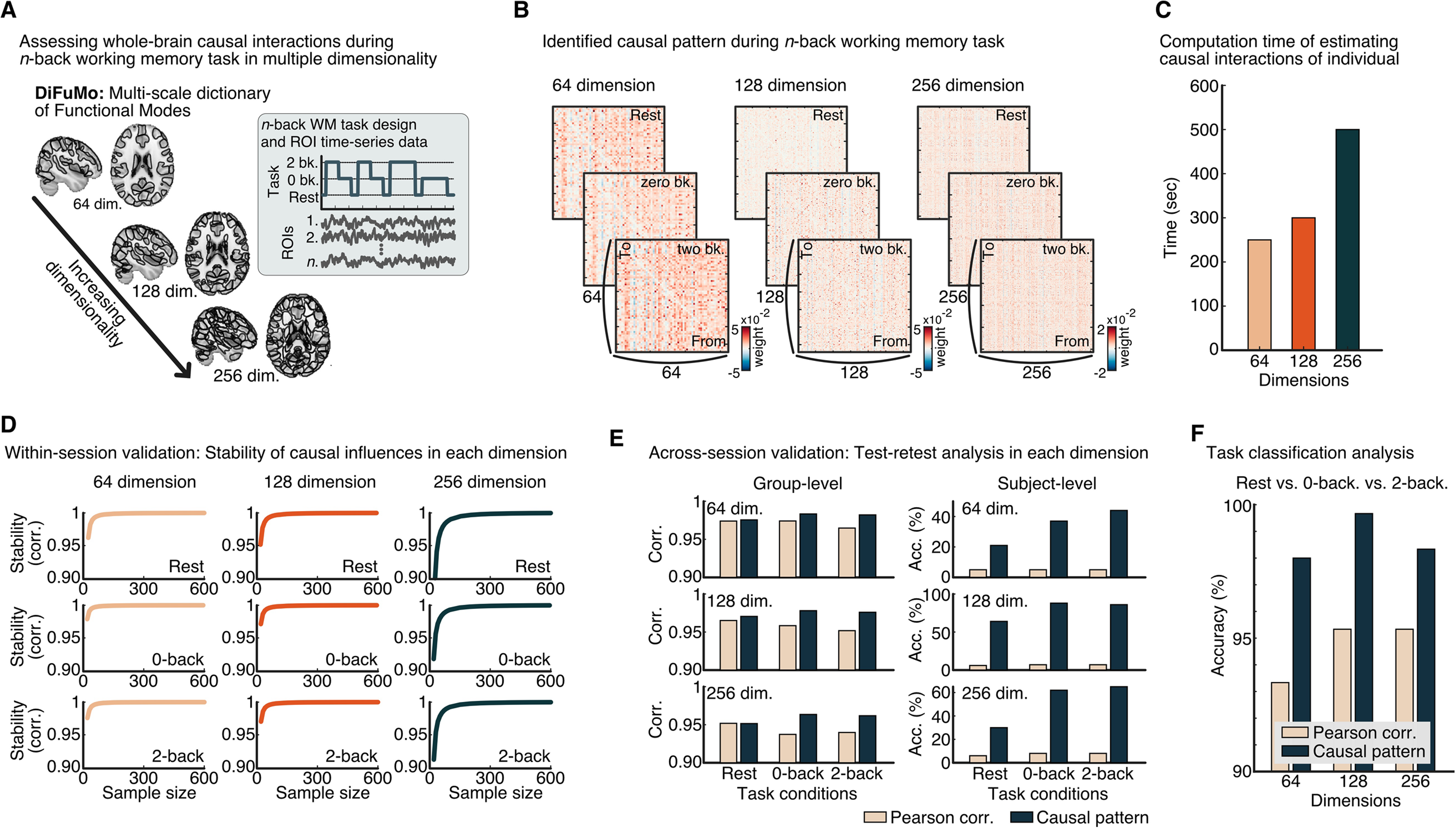
MDSI-AVI demonstrates robust scalability and reliability for whole-brain causal modeling using the DiFuMo atlas. **(A)** Multi-scale analysis approach across 64, 128, and 256 brain regions during n-back working memory tasks. **(B)** Significant causal interaction patterns identified across all spatial scales and task conditions. **(C)** Individual-level computational efficiency, with processing times scaling linearly from 250 to 500 seconds across increasing network sizes. **(D)** Stability analysis reveals high reliability (r = 0.90) with minimal sample sizes (N ≥ 20) across all conditions and scales. **(E)** Cross-session validation demonstrates excellent test-retest reliability (r > 0.95) and superior subject identification compared to Pearson correlation methods. **(F)** Task classification achieves >98% accuracy (*p* < 0.01, permutation test) in distinguishing cognitive conditions across all scales, substantially outperforming traditional connectivity approaches.

Stability analyses revealed that dynamic causal interaction patterns achieved a high level of stability (r=0.90, Pearson’s correlation) with subsample sizes of N=20 or more (**Figure 4D**) across all task conditions and dimensions. Group-level estimates of context-dependent brain-wide causal matrices were highly consistent across two scan sessions (r > 0.95; **Figure 4E, left**), and MDSI-AVI estimates outperformed Pearson-correlation-based features in subject prediction across all conditions and dimensions (**Figure 4E, right**).

Multivariate patterns of whole-brain dynamic causal interactions distinguished between task conditions (rest vs. 0-back vs. 2-back) with over 98% accuracy across all dimensions (*ps*<0.01, permutation test, **Figure 4F**) in left-out subjects, surpassing the predictive power of traditional Pearson correlation analyses.

These results demonstrate that MDSI-AVI reliably estimates dynamic causal interaction patterns associated with task conditions and exhibits high participant specificity, scalability, and accuracy in capturing brain-wide causal circuit dynamics during working memory.

### Whole-brain modeling of causal circuit dynamics using DiFuMo: consistency of MDSI-AVI estimates across different parcellation scales

To assess the similarity of estimates derived from multiple scales, we assigned each region in each scale to one of seven intrinsic functional systems ^61^ and constructed system-by-system functional causal matrices from each scale (**Figure 5A**). We then computed pairwise Pearson correlations of system-level causal patterns derived from different scales for each condition. This analysis revealed high correlations (r > 0.88) of system-by-system causal patterns derived from different scales (**Figure 5B**), indicating strong consistency in system-level causal interactions across scales. Additional analysis demonstrated that the correlation differences between task conditions are greater than those between scales, indicating that task context has a stronger influence on the estimated causal interactions than the choice of parcellation (**Supplementary Figure 5**). These findings suggest that MDSI-AVI produces highly consistent estimates of large-scale causal brain dynamics across different spatial resolutions, while also being sensitive enough to detect meaningful task-related changes in causal architecture.

**Figure 5.**
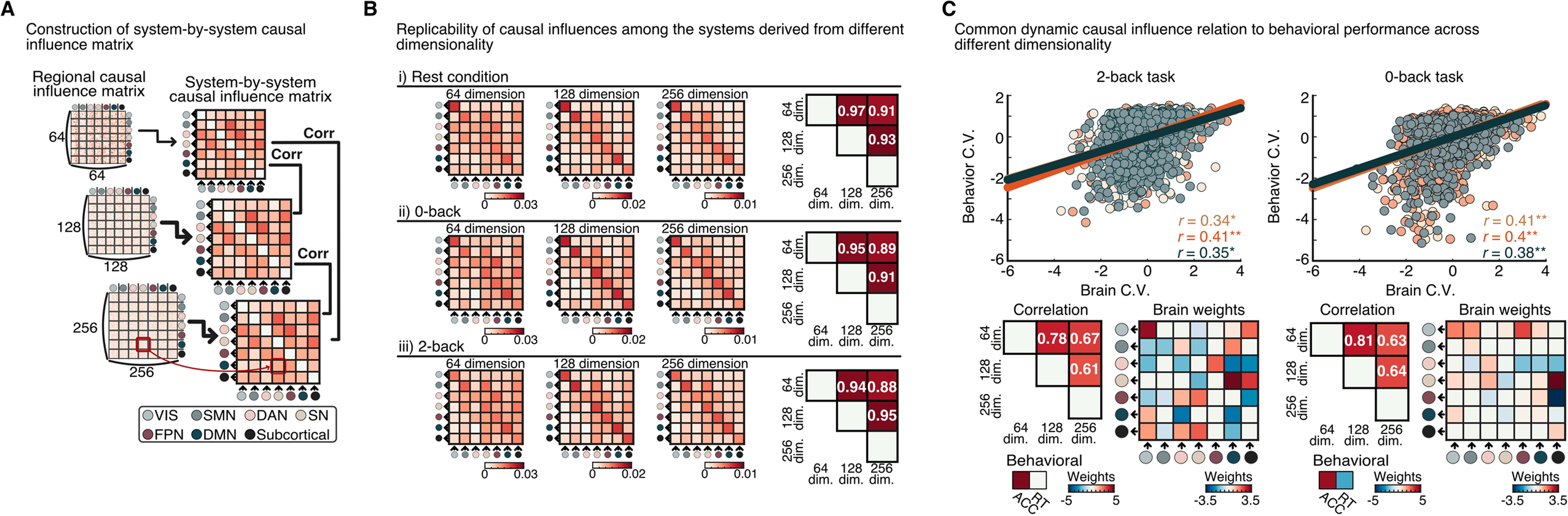
MDSI-AVI produces consistent network-level insights across multiple spatial scales. **(A)** Analysis strategy for aggregating regional causal interactions into system-level patterns across seven major brain networks. **(B)** Cross-scale consistency demonstrates high correlations (*r* > 0.88) of network-level causal patterns across different spatial resolutions, indicating robust multi-scale organization. **(C)** Brain-behavior relationships are preserved across scales. Canonical correlation analysis reveals consistent associations between network-level causal dynamics and working memory performance (accuracy and reaction time) in both task conditions, with Salience-Frontoparietal-Default Mode network interactions showing the strongest and most reproducible behavioral relevance.

We then performed Canonical Correlation Analysis (CCA) to determine the relationship between condition-dependent system-level causal patterns derived from each scale and behavioral measures (accuracy and reaction time) in each participant. Permutation testing revealed significant CCA components linking behavior scores and system-level causal weights derived from each scale in both 2-back (**Figure 5C, left**, FDR-corrected, *ps*<0.001) and 0-back conditions (**Figure 5C, right**, FDR-corrected, *ps*<0.05). The canonical causal weights across scales showed high correlations, with specific causal weights between DMN, SN, and FPN being the strongest and shared across scales.

These results demonstrate that MDSI-AVI can reliably estimate brain-wide causal dynamics, resulting in coherent causal system dynamics across different spatial scales. Moreover, the causal patterns derived from each scale inherit behaviorally relevant information, highlighting the power of MDSI-AVI in robustly extracting behaviorally meaningful causal dynamics from various scales.

### Generalizability to an independent anatomically-informed whole-brain atlas

To further validate the robustness and generalizability of MDSI-AVI, we sought to replicate our findings using an independent, anatomically-informed brain atlas. This approach allows us to assess whether our results are consistent across different parcellation schemes and to potentially gain more detailed insights into the anatomical substrates of the observed causal dynamics.

We employed the Brainnetome atlas, which provides a fine-grained parcellation based on both structural and functional connectivity profiles, closely corresponding with cytoarchitectonic features and the Brodmann atlas ^62^. This atlas offers 246 brain regions, providing a more detailed anatomical resolution compared to the DiFuMo atlas used in our previous analyses.

We estimated condition-dependent whole-brain causal interactions using this parcellation (**Figure 6A**).

**Figure 6.**
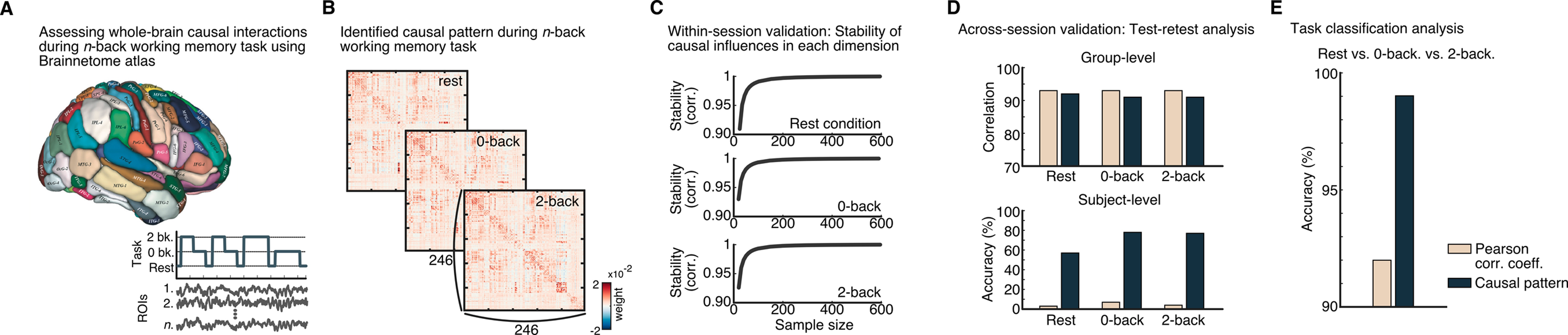
Independent validation using the anatomically-informed Brainnetome atlas confirms MDSI-AVI reliability. **(A)** Whole-brain causal analysis using the 246-region Brainnetome parcellation during n-back working memory tasks. **(B)** Robust causal interaction patterns identified across all task conditions with high statistical significance. **(C)** Stability analysis confirms high reliability with subsample sizes N ≥ 20, replicating findings from the DiFuMo atlas. **(D)** Cross-session validation demonstrates excellent group-level consistency (*r* > 0.9) and superior individual subject classification compared to correlation-based methods. **(E)** Task classification achieves >95% accuracy, confirming the generalizability of MDSI-AVI’s superior performance across different brain parcellation schemes and demonstrating robustness of the whole-brain causal modeling approach.

MDSI-AVI identified asymmetric connectivity patterns exhibiting significant dynamic causal interactions under rest, 0-back, and 2-back task conditions (*p*<0.01, FDR-corrected, two-sided paired t-test) (**Figure 6B**). Importantly, we observed high stability of MDSI-AVI estimates with subsample sizes of N=20 or more (**Figure 6C**) across all task conditions, replicating our findings from the DiFuMo atlas. Stability analyses demonstrated high group-level consistency (*r* > 0.9, **Figure 6D**) across sessions and superior subject classification accuracy compared to the Pearson-correlation-based approach. Furthermore, the causal patterns derived from the Brainnetome atlas distinguished different working memory task conditions (2-back vs. 0-back vs. rest) with remarkable accuracy (> 95%), again surpassing the predictive power of traditional Pearson correlation analyses (**Figure 6E**).

These results demonstrate the generalizability of the MDSI-AVI model, showing its ability to robustly and reliably estimate dynamic causal interaction patterns associated with task conditions across different brain parcellation schemes. The consistency of our findings between the DiFuMo and Brainnetome atlases strengthens the validity of our approach. Moreover, the use of the anatomically-informed Brainnetome atlas provides a foundation for more precise anatomical interpretation of the observed causal dynamics in subsequent analyses.

### Salience network as dominant causal outflow network in whole-brain dynamics

Previous research has suggested that the Salience Network (SN) plays a crucial role in network switching and cognitive control ^39,63–69^. However, these studies have been limited to examining interactions among a small number of brain regions. To comprehensively understand the SN’s role in working memory, it is essential to investigate its causal influences at the whole-brain level. We hypothesized that SN nodes would exhibit the highest increase in causal outflow during working memory tasks compared to the resting state, reflecting their proposed role in coordinating large-scale network dynamics.

To identify task-dependent changes in causal circuit dynamics during working memory, we contrasted the causal interactions of individual brain regions during the 2-back and 0-back tasks with those observed during the resting condition. This whole-brain approach allowed us to examine the SN’s influence on a broader scale, including its interactions with all nodes of multiple other key networks such as FPN and DMN.

Our analysis revealed heterogeneous changes in inter-regional causal circuit dynamics, with regions in the SN acting as dominant outflow nodes (**Figures 7A and 7B**). Specifically, MDSI-AVI identified a significant increase in causal interactions from SN nodes to FPN and DMN nodes during both the 2-back and 0-back tasks, compared to the resting baseline (*p* < 0.05, FDR-corrected, two-sided paired *t*-test). Notably, during the 2-back task, DMN nodes showed the strongest increase in inflow compared to rest (*p* < 0.05, FDR-corrected, two-sided paired t-test), suggesting a potential regulatory influence of the SN on DMN activity during higher cognitive load.

**Figure 7.**
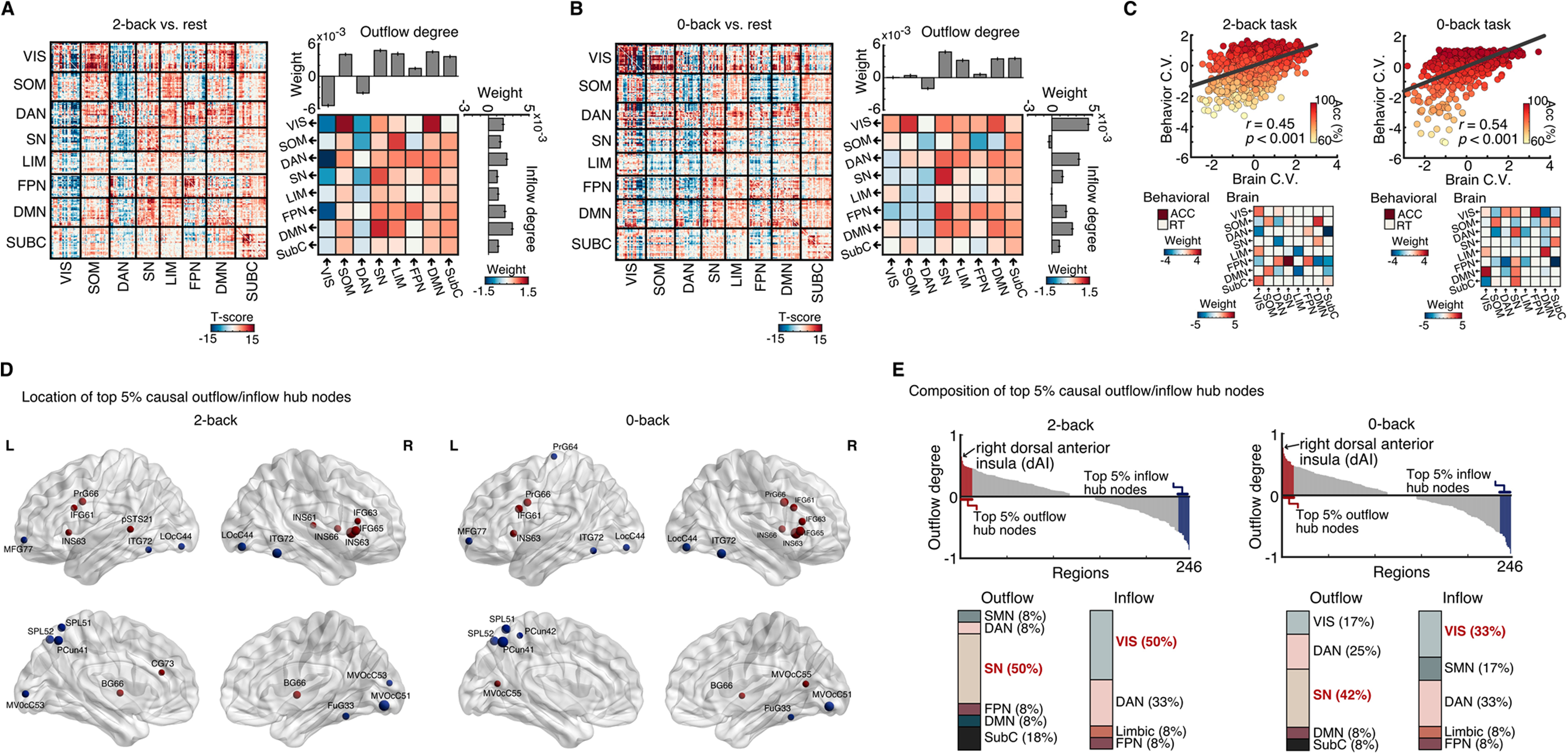
The Salience Network emerges as the dominant causal outflow hub during working memory tasks. **(A)** Task-dependent changes in 2-back working memory. Left: Region-level causal interaction changes compared to rest reveal asymmetric patterns across the brain. Right: Network-level analysis shows the Salience Network (SN) as the primary causal outflow source, with the Frontoparietal Network (FPN) and Default Mode Network (DMN) as major targets. **(B)** Task-dependent changes in 0-back working memory. Similar outflow dominance of the SN is observed, though with different target network patterns compared to 2-back conditions. **(C)** Causal dynamics predict working memory performance. Canonical correlation analysis demonstrates that SN→FPN and SN→DMN causal influences significantly correlate with behavioral performance in both 2-back (r = 0.45, p < 0.001) and 0-back (r = 0.54, p < 0.001) conditions. **(D-E)** Anatomical specificity of causal hubs. The right dorsal anterior insula node of the SN emerges as the strongest causal outflow hub across both conditions (p < 0.05, FDR-corrected), alongside bilateral inferior frontal gyrus. Major inflow hubs include left middle frontal gyrus, superior parietal lobule, left precuneus, and right fusiform gyrus, revealing a hierarchical organization of information flow during working memory.

To investigate the behavioral relevance of these dynamic causal interactions, we employed CCA. This analysis revealed significant relationships between dynamic causal weights and behavioral scores (accuracy and reaction time) in both the 2-back condition (*r* = 0.45, *p* < 0.001, **Figure 7C, left**) and the 0-back condition (*r* = 0.54, *p* < 0.001, **Figure 7C, right**). Notably, in the 0-back condition, the strongest weights were associated with connections from the SN to both the FPN and DMN, whereas in the 2-back condition, strong weights were primarily observed from the SN to the FPN. These findings highlight the behavioral relevance of dynamic causal interactions among the SN, FPN, and DMN.

### Anatomical specificity of causal hub nodes engaged during working memory

While network-level analyses provide valuable insights into large-scale brain dynamics, examining node-level specificity is crucial for pinpointing the exact anatomical loci responsible for coordinating these dynamics. Previous studies have suggested a key role for the anterior insula in network switching, but have often treated the insula as a homogeneous structure.

However, recent evidence suggests functional heterogeneity within the insula ^70^. By leveraging our whole-brain approach and the fine-grained Brainnetome atlas, we aimed to test the hypothesis that specific subdivisions of the insula, particularly the dorsal anterior insula (dAI), play a dominant causal role in network switching during working memory.

We evaluated the net causal influences of each node and determined causal signaling hubs during performance of the two working-memory tasks. Specifically, we identified the composition of the top 5% causal outflow and inflow hub nodes (**Figure 7D, E** and **Supplementary Table 2-5**). Our analysis revealed that the right dAI and bilateral inferior frontal gyrus were among the highest causal outflow hub nodes in both 2-back and 0-back conditions. Notably, the dAI in the right hemisphere showed the highest outflow degree in both conditions (*p* <0.05, FDR-corrected, two-sided paired t-test). In contrast, the left middle frontal gyrus, superior parietal lobule, left precuneus, and right fusiform gyrus were identified as top causal inflow hubs (**Figure 7D, E** and **Supplementary Table 2-5**).

These findings provide unprecedented anatomical specificity in identifying causal hub nodes during working memory tasks. The emergence of the right dorsal AI as the strongest causal outflow hub across both task conditions supports its hypothesized role in coordinating large-scale network dynamics. This specificity goes beyond previous studies that treated the AI as a homogeneous structure, highlighting the importance of examining brain function at a finer anatomical scale.

### Whole-brain network-level causal influences in human working memory

While identifying causal hub nodes provides valuable insights, understanding the nature of information flow in the brain is crucial for elucidating the mechanisms of cognitive control during working memory. Examining both direct and indirect causal influences provides a more comprehensive view of how key regions, such as the right dAI, may influence network dynamics. This approach can reveal how information potentially propagates through multiple pathways in the brain, offering insights into the neural processes involved in working memory.

Having identified the right dAI as the highest outflow hub node in both 2-back and 0-back conditions, we sought to examine how it influences cognitive systems during working memory. We analyzed both direct and indirect causal pathways originating from the right dAI to other brain networks.

In the 2-back condition, our analysis revealed a distinct pattern of causal influences (**Figure 8A**). The FPN received the highest direct influence from the right dAI (p<0.05, FDR-corrected, two-sided paired t-test). The DMN showed the highest indirect influence, mediated through the FPN (*p*<0.05, FDR-corrected, two-sided paired *t*-test).

**Figure 8.**
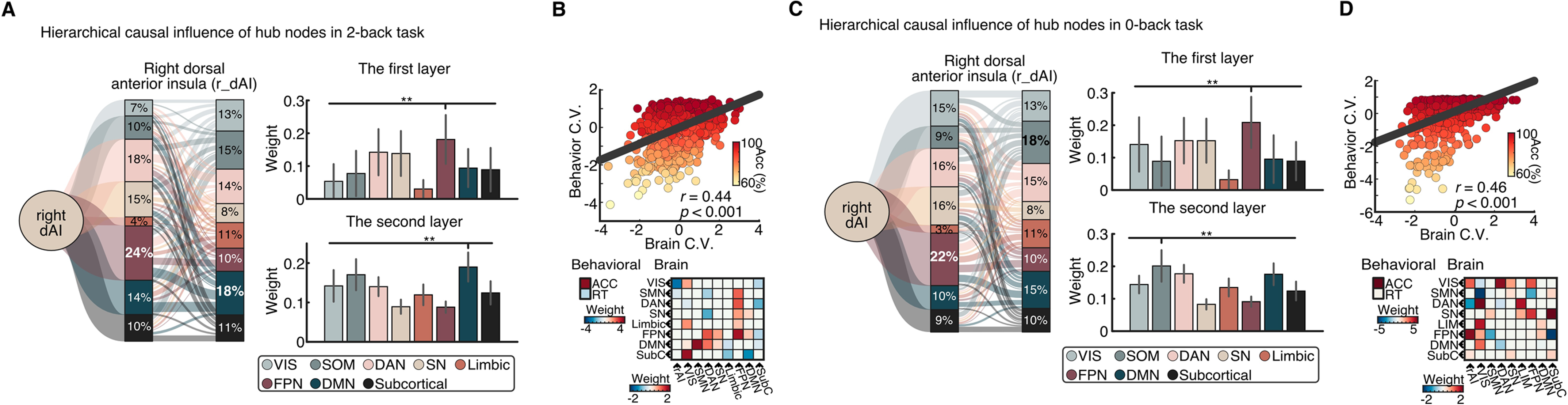
Load-dependent hierarchical information flow reveals task-specific causal pathways. **(A)** 2-back condition: Multi-stage causal cascade. The dorsal anterior insula (dAI) exerts strongest direct influence on the Frontoparietal Network (FPN), which in turn drives Default Mode Network (DMN) activity (*p* < 0.05, FDR-corrected), creating a dAI→FPN→DMN pathway. **(B)** Brain-behavior coupling in 2-back condition. These causal pathways significantly predict working memory performance (r = 0.44, p < 0.001). **(C)** 0-back condition: While dAI→FPN influence remains consistent, the FPN drives Sensorimotor Network (SOM) rather than DMN activity, creating a dAI→FPN→SOM pathway that reflects the simpler motor demands of this condition. **(D)** Brain-behavior coupling in 0-back condition. This alternative causal pathway also significantly predicts performance (r = 0.46, p < 0.001), demonstrating that different cognitive demands engage distinct downstream targets while maintaining consistent top-down control from the dAI.

Interestingly, the pattern differed in the 0-back condition (**Figure 8C**). While the FPN still received the highest direct influence from the rAI, the sensorimotor network showed the strongest indirect influence associated with the FPN (*p*<0.05, FDR-corrected, two-sided paired *t*-test). To assess the behavioral relevance of these influences, we conducted canonical correlation analyses with condition-dependent causal weights, which comprised both the direct and indirect causal pathways originating from the right dAI to other brain networks. These revealed significant relationships between direct and indirect causal influences from right dAI and behavioral performance in both the 2-back (*r* = 0.44, *p* < 0.001, **Figure 8B**) and 0-back (*r* = 0.46, *p* < 0.001, **Figure 8D**) conditions.

These findings provide novel insights into the nature of causal influences during working memory. The consistent primary influence of the rAI on the FPN across both task conditions supports the hypothesis that the rAI plays a crucial role in initiating cognitive control processes. However, the divergence in secondary influences between the 2-back (to DMN) and 0-back (to SOM) conditions suggests that different networks may be engaged based on task demands.

### Comparison with regression DCM model

Finally, we benchmarked MDSI-AVI against regression-based Dynamic Causal Modeling (rDCM), a widely used framework for whole-brain causal connectivity analysis ^71–73^ (**Supplementary Discussion**). Unlike MDSI-AVI, rDCM assumes a fixed canonical HRF, limiting its flexibility to capture regional and subject-specific hemodynamic variability.

We first evaluated model performance using simulated fMRI datasets with known ground-truth connectivity, assessing both connection strength estimation (Mean Absolute Percentage Error, MAPE) and binary edge detection (Area Under the ROC Curve, AUC). Despite the ground-truth data being generated using a DCM-based model, which inherently favors rDCM, MDSI-AVI consistently outperformed rDCM across diverse network configurations. This advantage was particularly pronounced for larger and more complex network structures (**Supplementary Table 1**), confirming that MDSI-AVI more accurately recovers both the topology and strength of directed connections.

We then compared performance on empirical HCP n-back working memory data across different parcellation scales using the DiFuMo and Brainnetome atlases (**Figure 9** and **Supplementary Figure 2 and 6**). At the group level, MDSI-AVI produced more stable causal matrices across scan sessions, indicating superior reliability (**Figure 9A**). At the individual level, MDSI-AVI achieved higher subject identification accuracy (**Figure 9B**). Task classification analysis further confirmed MDSI-AVI’s superior performance (**Figure 9C**). These findings highlight the advantages of MDSI-AVI’s flexible variational inference framework and underscore the critical importance of modeling regional hemodynamic heterogeneity for reliable, individualized estimates of causal brain dynamics.

**Figure 9.**
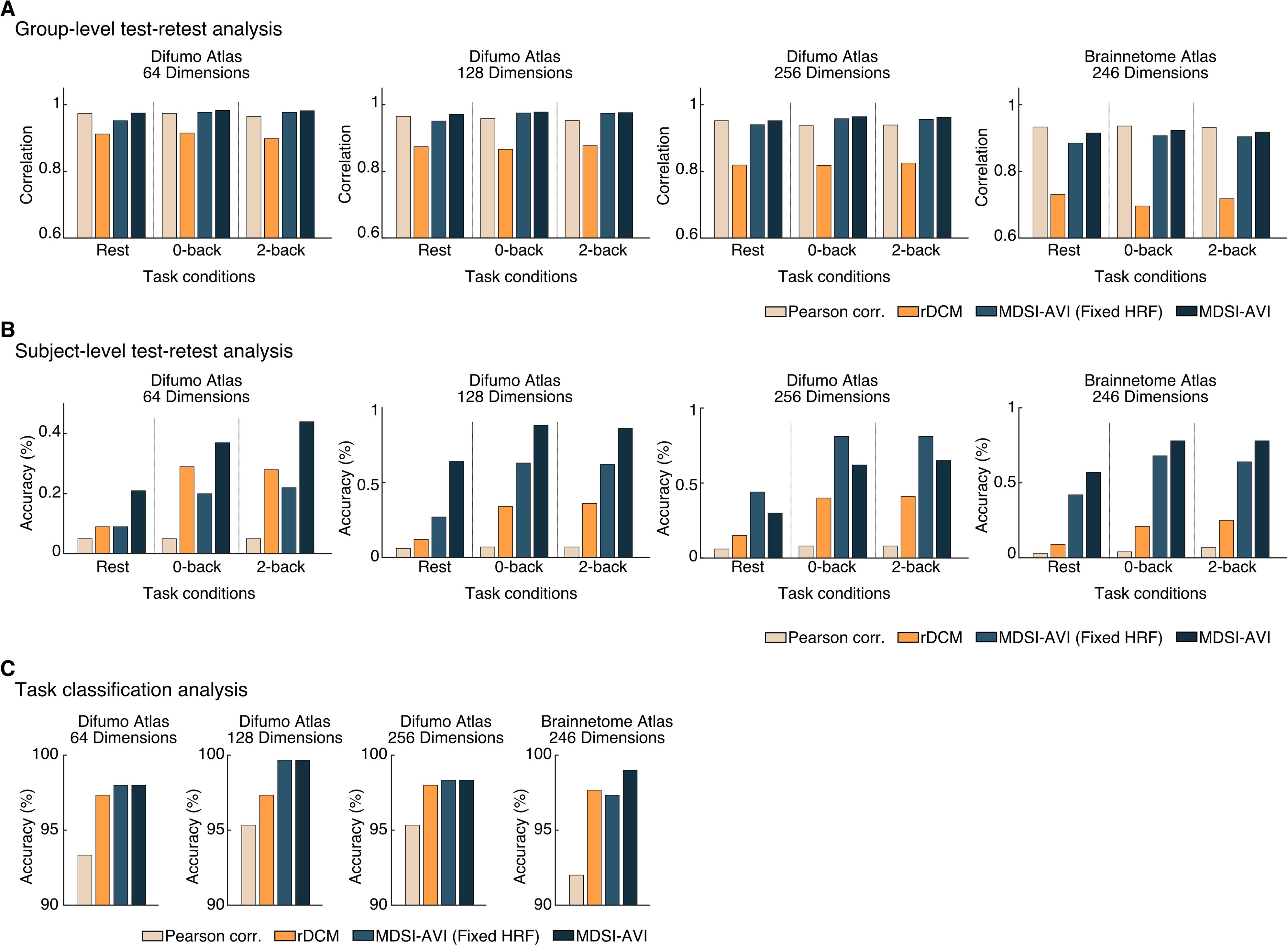
MDSI-AVI demonstrates superior performance across multiple validation criteria compared to established methods. **(A)** Test-retest reliability. MDSI-AVI achieves the highest group-level stability across scan sessions, tasks, and parcellation scales compared to Pearson correlation, rDCM, and fixed-HRF variants, indicating superior reliability for estimating causal brain dynamics. **(B)** Individual subject specificity. MDSI-AVI shows markedly superior subject identification accuracy, with the full model consistently outperforming its fixed-HRF variant. This demonstrates that modeling regional hemodynamic variability is crucial for capturing individualized brain dynamics and achieving reliable person-specific estimates. **(C)** Task sensitivity. MDSI-AVI achieves >98% accuracy in distinguishing cognitive conditions, substantially exceeding both rDCM and correlation-based approaches across all spatial scales. This superior task classification performance highlights MDSI-AVI’s enhanced sensitivity to functionally relevant changes in causal brain organization during cognitive tasks.

### Comparison with MDSI fixed HRF variant

To isolate the specific contribution of hemodynamic variability modeling, we compared MDSI-AVI to a variant that assumed a fixed canonical HRF across all regions and subjects while retaining the same hybrid variational inference strategy and model structure (see **Methods** for details). This controlled comparison enabled direct assessment of the benefits conferred by modeling HRF variability.

In simulated datasets, both models performed similarly, with MDSI-AVI showing slightly better average estimation accuracy (lower MAPE: 13.8% vs. 14.1%) and edge detection performance (higher AUC: 0.88 vs. 0.86). These gains, while small, support the theoretical advantage of incorporating HRF variability, particularly for larger networks (**Supplementary Table 1**).

Crucially, performance differences became more pronounced in empirical analyses, where MDSI-AVI consistently outperformed its fixed-HRF counterpart across all metrics (**Figure 9**). Most notably, subject-level test-retest reliability, assessed through subject identification accuracy, was substantially higher for MDSI-AVI across multiple parcellation granularities. These findings demonstrate that while both approaches perform comparably under controlled simulation conditions, explicitly modeling HRF variability provides clear advantages in real-world applications by more accurately capturing individualized neural dynamics and improving model reliability.

## DISCUSSION

Our study addresses several longstanding limitations in understanding whole-brain dynamic circuit mechanisms underlying cognition. We developed and validated MDSI-AVI, an innovative computational framework designed to capture asymmetric fMRI task-related directed interactions at the whole-brain level. This approach represents a major advance over prior studies that have relied on a small number of pre-specified regions of interest, potentially missing crucial brain-wide dynamics.

Application of MDSI-AVI to an n-back working memory task revealed behaviorally-relevant asymmetric causal interactions across the entire human brain. Notably, these dynamic causal interactions distinguished between different working memory loads (2-back vs. 0-back) with remarkable accuracy (>95%), surpassing the predictive power of traditional Pearson correlation analyses. Moreover, the strength of causal interactions robustly predicted memory performance. These findings were replicated across multiple atlases and different spatial parcellations, demonstrating the robustness of our approach.

Our whole-brain analysis identified the salience network as a dominant causal outflow network, with the dorsal anterior insula (dAI) emerging as a key hub. This finding provides unprecedented anatomical specificity in identifying causal nodes during working memory, going beyond previous studies that could not address anatomical specificity without testing multiple models. MDSI-AVI also allowed us to elucidate multi-stage hierarchical causal chains for the first time, revealing a strong indirect causal influence from the SN to the FPN, with unique load-dependent downstream effects on the DMN. The strength of these hierarchical interactions increased with cognitive load and predicted working memory performance.

Our state-space modeling approach provides a powerful and flexible toolbox for probing neural mechanisms underlying cognition and behavior across the entire brain, setting the stage for more comprehensive investigations of various cognitive processes and their disruptions in neurological and psychiatric conditions.

### Addressing inference limitations through hybrid variational inference

Our approach represents a significant methodological advance in the emerging paradigm toward simulation-based inference. Variational inference through model-based simulations is rapidly becoming a transformative approach across scientific disciplines ^74,75^, offering principled ways to handle complex, high-dimensional inference problems that were previously intractable ^74–78^. Our method addresses fundamental limitations of classical variational inference—particularly mode collapse and over-precision—through two key innovations: (1) a hybrid variational inference (h-VI) strategy and (2) explicit modeling of hemodynamic response function (HRF) variability across brain regions and individuals.

First, we leveraged a hybrid two-step optimization framework that combines amortized inference with variational optimization (**Figure 1B**). In the first step, we train an encoder and a normalizing flow to learn the distributions of the hyper-parameters, latent noise (**q**), observed noises (**r**), and hemodynamic parameters (**H**) given synthetic BOLD signals. This amortized inference is crucial because the mapping from latent variables to BOLD is many-to-one—i.e., multiple configurations can yield similar BOLD signals. Learning these distributions enables us to marginalize their uncertainty efficiently in downstream inference. In the second step, we perform subject-specific inference of the latent neuronal activity (**X**) and the directed connectivity matrix (**A**) by minimizing the reverse Kullback-Leibler divergence. Compared to rDCM, which uses an iterative scheme to minimize the r-KL, our model takes advantage of automatic differentiation to achieve this while also automatically marginalizing the uncertainty of the noise and the HRF for each region, enabling both scalability to large brain networks and individual-specific modeling.

Second, incorporating HRF variability into the generative model introduces necessary flexibility for disentangling neural activity from hemodynamic effects. Unlike models with fixed canonical HRFs, our approach allows region-and subject-specific variations in HRF dynamics, thereby reducing confounds that could bias the estimation of neural interactions. This added expressiveness not only enhances the physiological plausibility of the model but also improves identifiability of latent neural causes, leading to more accurate and generalizable causal inference. Together, these methodological advances expand the posterior distribution accessible to the model, reduce inference artifacts, and improve the reliability of subject-level directed connectivity estimates—particularly in the presence of inter-individual variability in neurovascular coupling.

### Validation, scalability and computational efficiency

MDSI-AVI represents a significant leap forward in analyzing whole-brain causal dynamics. A key innovation is its ability to scale to the entire brain, overcoming limitations of previous methods restricted to a few pre-specified regions ^7,25,26,79^. We successfully applied MDSI-AVI to brain atlases with 64, 128, and 256 regions, demonstrating robust performance across different levels of granularity.

Importantly, we validated MDSI-AVI using both simulated data and optogenetic stimulation experiments, providing complementary forms of validation. In simulated data, where ground truth is known, MDSI-AVI achieved high AUC and low MAPE values, demonstrating its accuracy in recovering true causal interactions. The simulations, ranging from simple networks to complex macaque connectome-based models, validated the method’s performance across different scales and complexities. Optogenetic stimulation experiments provided a crucial biological validation, allowing us to test MDSI-AVI’s ability to detect known causal influences in real brain data. This combination of *in silico* and *in vivo* validations establishes a strong foundation for the reliability of MDSI-AVI in real-world applications.

A critical advance is MDSI-AVI’s ability to account for hemodynamic response function variability across a large number of brain regions. This feature allows for more accurate representation of latent neural activity and avoids biases in causal estimates that can arise from assuming a uniform HRF (**Figure 3**). An innovation of our model is that our hybrid approach separates the parameters into two groups treated with different inference methods, allowing for robust marginalization of HRF coefficient uncertainty and scalable inference of thousands of regional causal interactions. This represents a significant advance over previous methods that assume a constant HRF across brain regions, which can lead to spurious or missed interactions ^80,81^. Our approach thus provides a more principled way to infer brain-wide interactions in fMRI data, enhancing the reliability and validity of our findings. This improvement is essential for advancing the field of cognitive neuroscience, as it ensures that the inferred interactions more accurately reflect the underlying neural dynamics rather than hemodynamic response function variability.

The computational efficiency of MDSI-AVI is its another major advantage. Unlike previous methods, our approach converged rapidly even for 256 regions of interest (**Figure 4C**). This efficiency represents a significant advance, as it allows for more extensive and in-depth exploration of whole-brain circuit dynamics without prohibitive time constraints. Furthermore, MDSI-AVI reliably identified brain-behavior relationships across each scale (**Figure 5C**). This scalability is crucial for future applications and research, allowing for broad brain-wide investigations unconstrained by the challenges of brain region selection procedures and model choice.

Finally, we demonstrated the reliability and reproducibility of our findings across different atlases, parcellation schemes, and subsamples. Stability analyses revealed that estimation of dynamic causal interaction patterns were highly stable across all task conditions and dimensionalities (**Figures 4, 6**). These analyses underscore the robustness of our approach and provide unique insights into reproducible dynamical systems-based mechanisms of human working memory. This robust cross-validation addresses critical concerns about replicability in neuroscience ^29^, providing a foundation for future studies.

These advances collectively enable a more accurate, comprehensive, and efficient analysis of whole-brain causal dynamics, opening new avenues for understanding complex cognitive processes and their neural underpinnings.

### Salience network as a causal hub in working memory at the whole-brain level

Previous research has suggested that the Salience Network (SN) plays a crucial role in network switching and cognitive control ^39,63–69^. However, these studies have been limited by their focus on interactions among a small number of pre-specified brain regions, potentially missing important whole-brain dynamics and introducing biases in the interpretation of the SN’s role. This limitation is particularly problematic when studying a network hypothesized to coordinate global brain activity, as the SN is hypothesized to do ^70,82^.

To comprehensively understand the SN’s role in working memory, it is essential to investigate its causal influences at the whole-brain level and contrast it with other networks which are also sampled brain-wide. This approach allows us to avoid selection bias in regions of interest, capture potential long-range interactions that might be missed in small-scale studies, and understand the SN’s influence in the context of the entire brain’s functional architecture. We hypothesized that SN nodes would exhibit the highest increase in causal outflow during working memory tasks compared to the resting state, reflecting their proposed role in coordinating large-scale network dynamics. Our whole-brain analysis using MDSI-AVI allowed us to test this hypothesis without the constraints of pre-specified ROIs.

Our findings provide unprecedented whole-brain evidence supporting the hypothesis that the SN plays a crucial role in coordinating large-scale network dynamics during working memory tasks. Specifically, we observed increased causal outflow from SN nodes to Frontoparietal Network (FPN) and Default Mode Network (DMN) regions. This suggests that the SN may aid in directing the engagement and disengagement of these networks based on task demands, a function that could not be fully captured in previous small-scale studies.

Importantly, the strong correlation between these causal interactions and behavioral performance underscores the functional significance of SN-mediated network switching in successful working memory processing. This brain-behavior relationship provides critical validation of the SN’s proposed role, linking its whole-brain causal influences directly to cognitive performance.

The high causal influence of the SN across the entire brain provides strong evidence that it may indeed act as a central hub for the flexible reconfiguration of brain networks in response to changing task demands ^7,43,44^. This finding extends previous theories about the SN’s role by demonstrating its global influence, rather than just its interactions with a few select regions.

Together, our findings not only confirm but significantly extend previous theories about the SN’s role in cognitive control. By overcoming the limitations of small-scale studies, we provide a more nuanced and complete understanding of how the SN may trigger brain-wide dynamics to support working memory function.

### Anatomical specificity of causal outflow and inflow hubs at the whole-brain level

A key scientific finding of our study is the identification of the dorsal anterior insula (dAI), a key node within the Salience Network, as a critical outflow hub at the whole-brain level. While previous research has implicated the dAI in a wide range of cognitive control processes ^7,43,44,70^, the use of a few pre-defined regions of interest has precluded a comprehensive understanding of its role within the broader context of whole-brain dynamics. Our approach allowed us to overcome this limitation, providing unprecedented anatomical specificity in identifying causal hub nodes during working memory tasks.

The emergence of the right dAI as the strongest causal outflow hub across both high and low working memory load conditions supports its hypothesized role in coordinating large-scale network dynamics. This finding aligns with the proposal that the dAI’s high causal outflow reflects its role in detecting salient stimuli, reallocating cognitive resources, and initiating control signals to other brain regions ^44,70^. Interestingly, the consistency of the dAI’s outflow across different working memory loads suggests that its role in salience detection and control initiation may be more general and not specific to working memory demands.

Our whole-brain, node-level approach goes beyond previous studies that treated the anterior insula as a homogeneous structure, highlighting the importance of examining brain function at a finer anatomical scale. The distinct patterns of causal outflow and inflow hubs provide insights into the nature of information flow during working memory tasks. The strong outflow from the right dAI and inferior frontal gyrus suggests their role in top-down control, while the inflow to FPN regions like the middle frontal gyrus and superior parietal lobule may reflect their involvement in maintaining and manipulating information. The contrasting patterns of causal influence observed in key nodes of the SN and FPN provide new insights into the functional roles of these networks in cognitive control and working memory. Specifically, the dominant outflow from SN nodes, particularly the dAI, coupled with the predominant inflow to FPN nodes, suggests a hierarchical organization of information processing. This asymmetry in causal influence aligns with theories proposing that the SN acts as an initial filter for salient information, subsequently engaging the FPN for more detailed processing and manipulation of task-relevant stimuli. Such a dynamic interplay between these networks could explain their consistent involvement across various cognitive control and working memory tasks ^65,83–85^, highlighting a general mechanism for coordinating brain-wide activity in response to cognitive demands.

Crucially, our findings converge on and significantly extend previous research on working memory networks. While prior studies were constrained to analyzing limited sets of brain regions associated within the SN, FPN and DMN, due to computational limitations. Our whole-brain approach reveals processes that operate within a broader neural architecture, addressing critical gaps in the literature that arose from the lack of multivariate models capable of capturing brain-wide interactions. By moving beyond region-specific studies to a comprehensive whole-brain perspective, we demonstrate how these networks dynamically interact across the entire brain to support complex cognitive functions^65,83–85^.

The identification of these specific anatomical loci as causal hubs has significant implications for understanding the neural basis of cognitive control and for developing targeted interventions. These results could inform the selection of stimulation sites for non-invasive brain stimulation techniques aimed at enhancing cognitive control in individuals with working memory impairments. The right dAI, in particular, emerges as a promising target for such interventions.

Our findings not only corroborate previous research on the importance of the dAI in cognitive control but also refine our understanding of its role by pinpointing specific subdivisions. This sets the stage for more targeted investigations of the neural mechanisms underlying working memory and cognitive control more broadly, potentially leading to more effective therapeutic strategies for cognitive disorders.

### Indirect causal information flow in working memory

Our study revealed distinct patterns of network interactions during working memory tasks, providing novel insights into potential roles of key networks in cognitive control. By examining both direct and indirect causal pathways, we uncovered patterns of information flow that go beyond previous studies limited to direct interactions between a few pre-specified regions.

Here, we define direct pathways as causal influences from one network to another, and indirect pathways as influences that may involve intermediary networks.

In the 2-back condition, we observed that the right dAI of the SN exerted a strong direct influence on the FPN, which in turn showed a strong influence on the DMN. Interestingly, in the 0-back condition, while the FPN still received a strong direct influence from the dAI, the FPN showed a stronger influence on the Sensorimotor Network (SMN) rather than the DMN.

These distinct patterns of interactions may reflect different cognitive processes involved in the 2-back and 0-back tasks. Both tasks require sustained attention to encode task-relevant features from external stimuli, which could be supported by interaction between the SN and FPN. However, the divergence in the second stage might reflect task-specific demands: the 0-back task may require a more direct motor response, while the 2-back task involves more complex processes of comparison and working memory updating.

The strength of indirect network interactions from the FPN to the DMN increased with working memory load, indicating enhanced signal flow under cognitively demanding conditions. This aligns with the idea that higher cognitive loads require more persistent and robust neural signaling ^86–88^. The strong correlation between these causal dynamics and behavioral performance underscores their functional significance, suggesting that effective working memory relies on the coordinated information flow between networks, not just the activation of individual regions.

Our ability to examine these interactions at the whole-brain level represents an advance over previous methods, which were limited in their scope. This approach extends our understanding beyond simple hub models of brain organization, revealing potential multi-stage propagation of causal influences. However, it is crucial to note that our fMRI-based method cannot directly infer the temporal sequence of these interactions. Further research using methods with higher temporal resolution, such as intracranial EEG, is needed to test the timing and sequence of these interactions. Nonetheless, our findings provide a framework for generating hypotheses about how the brain implements cognitive control, potentially informing more sophisticated models of working memory and cognitive processing.

This perspective opens new avenues for investigating cognitive dysfunctions. By examining alterations in these network interactions, we may gain insights into the specific processes affected in various cognitive disorders, potentially leading to more targeted therapeutic interventions.

## Conclusion

Our study presents a powerful and flexible computational framework, MDSI-AVI, for analyzing whole-brain causal interactions underlying human cognition. This novel approach addresses critical challenges in whole-brain causal modeling—such as scalability, HRF variability, and contextual modulation—opening new avenues for understanding the neural basis of working memory and cognitive control.

A key advance of our whole-brain methodology is its ability to uncover both direct and indirect, multi-stage information flow across the entire brain. This capability revealed further insights into how the Salience Network, particularly the right dAI, influences load-dependent activity in other networks like the Frontoparietal and Default Mode networks during working memory tasks. The identification of these hierarchical causal chains provides a richer framework for understanding how the brain implements cognitive control, moving beyond simple hub models to a more sophisticated, multi-stage propagation of causal influences.

The scalability, speed, and accuracy of MDSI-AVI help pave the way for future research, advancing our understanding of brain function and brain-behavior relationships in the context of causal asymmetric directed interactions. By bridging the gap between localized and distributed perspectives on brain function, our approach offers a more complete and integrative understanding of how the brain orchestrates cognition and behavior.

While our approach represents a significant advance, future work should focus on extending this method to other cognitive domains and clinical populations. Integration with other neuroimaging modalities could provide a multi-scale understanding of brain dynamics.

Particularly promising is the potential to examine disruptions in hierarchical causal chains in various cognitive disorders, potentially gaining insights into the specific stages at which information processing breaks down. This could lead to more targeted therapeutic interventions, tailored to address specific disruptions in the causal flow of information in neurological and psychiatric disorders.

## METHODS

### MDSI generative model

Here, we briefly describe the MDS generative model. We hypothesize that the observed blood oxygenation level-dependent (BOLD) signal is generated by the convolution of some latent neural activation with the hemodynamic response function (HRF). We assume this coupling between HRF and the latent activation to be linear. The MDS generative model can be summarized via the following set of equations:

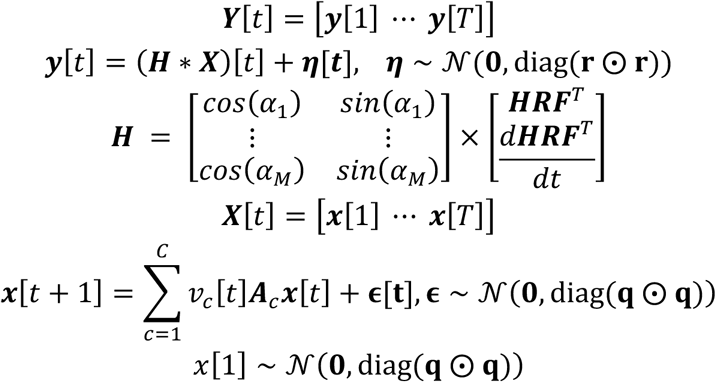

***Y*** ∈ ℝ*^M^*^×*T*^is the observed BOLD signal measured across *M* regions and *T* time points. The BOLD signal ***Y*** is modeled as the convolution of the latent neural activations ***X*** ∈ ℝ*^M^*^×*T*^with an HRF kernel ***H*** ∈ ℝ*^M^*^×*K*^ plus a zero-mean Gaussian noise ***η*** ∼ *N*(**0**, diag(**r** ⨀ **r**)), where *K* denotes the temporal duration of the HRF and diag(**r** ⨀ **r**) ∈ ℝ*^M^*^×*M*^ is the observation noise covariance matrix. Both the HRF kernel ***H*** and the observation noise ***η*** are region-specific: Each region *m* ∈ {1, *M*} is associated with a noise variance *σm*^2^ and an angle 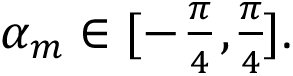 Additionally, Observation noise vector, ***η***, at time instance 1, 2,…,*T* are assumed to be identical and independently distributed (iid). The angle *α_m_* defines the HRF kernel as *cos* (*α_m_*) ***HRF*** +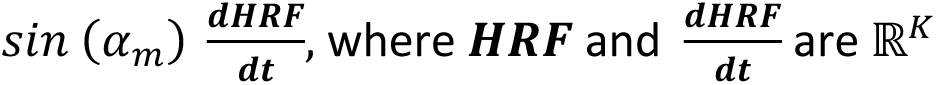 vectors corresponding respectively to the canonical HRF computed by the ‘*spm_hrf’* function in SPM12^89^ and its time derivatives.

***X*** ∈ ℝ*^M^*^×*T*^ represents the latent neural activations across *M* regions and *T* time points.

***x***[*t* + 1] ∈ ℝ*^M^*^×1^ denotes the latent neural activations at time *t* + 1 across *M* regions. At a given instance *t* + 1, the activations ***x***[*t* + 1] ∈ ℝ*^M^*^×1^ are obtained by a linear multiplication of modulatory inputs *v_c_*[*t*] ∈ {0, 1}, a region-to-region coupling matrix ***A****_c_* ∈ ℝ*^M^*^×*M*^, and the latent signal at previous time point ***x***[*t*] ∈ ℝ*^M^*^×1^, summed across experimental conditions *c* ∈ {1, *C*}, where *C* denotes the number of experimental conditions. *v_c_*[*t*] is the indicator function for condition *c*, which follows *v_c_*[*t*] = 1 if, and only if, *c* is the condition the subject is undergoing at time *t* ∈ {1, *T*}. The set of functions *v_c_*[*t*] for all *c* is defined as *v*[*t*]. We defined ***A****_c_* as the coupling matrix at condition *c*, denoting ***A*** the set of coupling matrices for all *c*. As such, we modeled the temporal evolution of latent neural signals in a condition-specific manner.

Additionally, we consider the addition of a zero-mean Gaussian noise **ε** ∼ N(**0**, diag(**q** ⨀ **q**)), where diag(**q** ⨀ **q**) ∈ ℝ*^M^*^×*M*^ denotes the state noise covariance matrix, modeling region-specific variability in latent neural dynamics. Additionally, state noise vector, **ε**, at time instance 1, 2,…, *T* are assumed to be identical and independently distributed (iid).

The quantity of interest of our analysis are the region-to-region coupling matrix ***A****_c_*. The element ***A****_c_*[*i*, *j*] represents how the previous latent signal in region *j* influences the current latent signal in region *i* under experimental condition *c*. Our goal is to infer the ***A*** ∈ ℝ*^C^*^×*M*×*M*^, from the observed BOLD signal **Y**. We hypothesize that **A** varies across conditions, deriving a matrix *A_c_* ∈ ℝ*^M^*^×*M*^that represents region-to-region coupling under each experimental condition.

### Hybrid Variational Bayes inference

Here we briefly describe our inference method. A more comprehensive treatment of the general framework of hybrid-VI can be found in ^90^. Following the Bayesian inference formalism, we estimated the posterior distribution *p*(***q***, ***r***, ***H***, ***X***, ***A***|***Y***, *v*). To do so, we use a variational Bayes method, also called Variational Inference (VI). We define a parametric distribution *q*(***q***, ***r***, ***H***, ***X***, ***A***) and optimize it to approximate the unknown posterior as closely as possible. For optimization, we employ the automatic differentiation variational inference (ADVI) framework ^91^, leveraging the reparameterization technique together with the automatic differentiation libraries and optimizers ^92^.

The term “hybrid” refers to our separation of the inferred latent parameters ***q***, ***r***, ***H***, ***X***, ***A*** into two groups, each associated with different training strategies and losses *L* minimized during training. We denote the first group as the hyper-parameters (HP), which includes the noise levels ***q***, ***r*** and the HRF kernel ***H***. A key challenge with those parameters is the existence of several disjoint sets of solutions that can explain the observed signal ***Y***. For instance, several combinations of region-specific HRFs and latent signals may yield equivalent reconstructions of ***Y***. Most of the conventional inference approaches struggle with those “multi-modal” scenarios, yielding a solution while ignoring equally plausible solutions—a phenomenon known as “*mode collapse*”. Mode collapse can result in underestimation of uncertainty and biased parameter estimation. To circumvent this issue, we train an amortized estimator *q_re_*_4*ion*_(*q_m_*, *r_m_*, *α_m_*; *f*(***y****_m_*)) that estimates a region’s noise level and HRF parameters given an encoding of the region’s BOLD signal ***y****_m_* ∈ *R^T^*. The encoder, represented by *f*, is implemented with a time convolutional neuronal network. The distribution *q_re_*_4*ion*_ modeled using a Masked Autoregressive Flow, is trained over a large dataset of synthetic examples generated from the model in Equation (1). The loss function used is: *L_HP_*_(*re*4*ion*)_ = *E_qm_*_,*r*_*_m_*_,*α*_*_m_*_,***y***_***_m_***_∼ *p*_(−*log q_re_*_4*ion*_(*q_m_*, *r_m_*, *α_m_*; *f*(***y****_m_*))), which (up to constants) is equivalent to the forward KL-divergence between the variational approximation and the true posterior. This training method prevents mode collapse. By considering all the regions independently, we obtain the HP estimator:

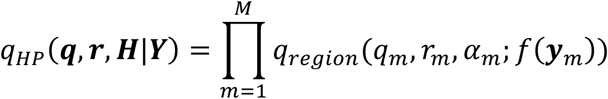

We denote the second parameter group as the parameters (P), including latent signal *X* and the coupling matrices *A_c_*′*s*. The main issue with those parameters is their large dimensionality. To ensure scalability to a large number of regions, we train the estimator *q_H_* to maximize the evidence lower bound (ELBO):

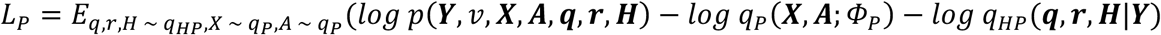

where *p* denotes the joint distribution defined by the generative model in Equation (1). *q_P_*, the P estimator, is a Gaussian mean-field with diagonal covariance, whose mean and variances, represented by *Φ_P_*, are the values of interest. The *q_HP_*, the HP estimator, is *not* trained during this second phase. Injecting the pre-trained *q_HP_* forces *q_P_* to consider all the possible HRFs that could explain the observed BOLD signal, preventing mode collapse for the latent signal ***X*** and in turn for the coupling matrix ***A***. Our hybrid strategy ensures fast and scalable inference while avoiding uncertainty underestimation or biased parameter estimation.

### Simulated fMRI dataset

We used several benchmarks of simulated fMRI data with known ground truth connectivity from Sanchez-Romero *et al.* ^58^. In general, this dataset included two groups of networks, one consisting of 9 simple small-scale synthetic graphs and one consisting of two graphs extracted from the macaque connectome. From the latter group, we used the smallest (‘Small-Degree Macaque’) and the largest (‘Full Macaque’) networks. The details of generating BOLD signals from each graph are detailed in Sanchez-Romero *et al.* ^58^. In brief, the same simulation procedure was used for simple and macaque-based graphs, where the authors used the model proposed in Smith *et al.* ^93^ which is itself based on the dynamic causal modeling (DCM) architecture ^36^. Underlying neural dynamics are simulated using the linear differential equation 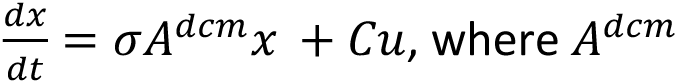 denotes the ground-truth connectivity. In this dataset, all connections between different regions are positive values. To simulate resting-state data, the **u** input was modeled using a Poisson process for each of the regions (*C* = *I*). The neuronal signals *x* were then passed through the Balloon-Windkessel model ^94–97^ to obtain simulated BOLD data. For each network configuration, 60 individual datasets were generated with introducing controlled variation in connection magnitudes and node-wise hemodynamic response functions (HRFs) with delays sampled from a Gaussian distribution (SD = 0.5 sec)

### AUC and MAPE measurements

To evaluate the directed couplings obtained from the simulated fMRI dataset two metrics were used: area under the receiver operator curve (AUC) and mean absolute percentual error (MAPE). For the AUC, the directed couplings were grouped by taking the absolute value of the population mean and dividing it by the population standard deviation for each element of the matrix (i.e., z-scored, excluding the diagonal). This population average was compared against the ground-truth connectivity *A^dcm^*, which was binarized assuming one for the existence of a connection and zero otherwise (also excluding the diagonal). This grouped-evaluation was motivated because, in this dataset, the position of the connections remains the same for all samples of a given network and only positive connections exist. The AUC was calculated using the popular Python library scikit-learn ^98^. For each subject and each element of the connectivity matrix, the MAPE was obtained by getting the absolute difference between the inferred value and the ground-truth connection divided by the ground-truth. To avoid division by zero, only true-connections were considered. The MAPE was averaged across links and subjects.

### Optogenetic stimulation dataset

Opto-fMRI data, including five adult female Sprague-Dawley rats (250–350 g; Charles River Laboratories, Wilmington, MA), were acquired. Two rats were excluded because one did not respond to optical stimulation and the second had movement-related artifacts. Of the final three rats included in this study, one was imaged at University of California, Los Angeles (UCLA) and two at Stanford University using identical imaging protocols. During surgery, M1 was targeted and injected with an adeno-associated virus expressing a ChR2-EYFP fusion protein using coordinates −2.7 mm anteroposterior (AP), +3.0 mm mediolateral (ML) right hemisphere,−2.0 mm and −2.5 mm dorsoventral (DV). Additional surgical procedures and details can be found in previous publications ^99,100^.

Experiments were conducted 3 weeks after virus injection for optimal ChR2 expression. The fMRI scans were performed on a 7T small animal MRI system (UCLA: Brucker Biospec, Stanford: Magnex Scientific). All scans used a 39 mm outer diameter and 25 mm inner diameter custom-designed transmit/receive single-loop surface coil. During the fMRI experiment, animals were artificially ventilated under light anesthesia with a mixture of O2 (35%), N2O (63.5%), isoflurane (1.2–1.5%) and CO2 (3–4%). A block-designed fMRI stimulation scheme consisting of six ON–OFF cycles at 20 s ON and 40 s OFF for a total of 6 min was used. During the ON cycles, optical stimulation was delivered at 20 Hz, with a 5 ms pulse duration. The data were acquired using an interleaved spiral readout Gradient Recalled Echo BOLD sequence with 0.5 mm slice thickness and 23 slices. In-plane field of view was designed to be 35×35 mm^2^ and in-plane spatial resolution was 0.5×0.5 mm^2^. A sliding window reconstruction was then performed to reconstruct the data into 128×128×23 matrix-size, 750 ms temporal resolution images.

After reconstruction, subject head motion was corrected by the inverse Gauss-Newton motion correction algorithm and 4D fMRI data was analyzed with statistical parameter mapping using the general linear model with five gamma basis. These five gamma-shaped basis functions, implemented in SPM12^89^, were used to flexibly model the hemodynamic response. By capturing potential variability in the timing and shape of the BOLD response to stimulation, they allow for a more accurate fit than a single canonical HRF. An F-test was then conducted and active voxels were selected as those with corresponding Bonferroni-corrected p-values < 0.05. The ROIs were manually selected based on a standard digital rat brain atlas ^101^.

### HCP n-back working memory dataset

HCP data from 728 individuals (age: 22–36 years old, 413 female/324 male) were selected from a total of 1200 subjects based on the following criteria: (1) participant had complete n-back task behavioral and fMRI data; (2) range of head motion in any translational and rotational direction <1 voxel; (3) average scan-to-scan head motion <0.2 mm; (4) accuracies in 0-back and 2-back conditions >50%; and (5) criterion (1)–(4) met in both sessions separately.

### HCP n-back working memory task

The HCP n-back working-memory task combines the category-specific representation task and the n-back working-memory task in a single-task paradigm. Subjects were presented with blocks of trials that consisted of pictures of faces, places, tools, and body parts. Within each session, the four different stimulus types were presented in separate blocks. Furthermore, within each session, half of the blocks are 2-back working-memory tasks and half are 0-back working-memory tasks. In the 2-back working-memory task blocks, subjects were requested to determine whether the current stimulus matches the stimulus in two presentations of stimuli prior within the same block. In the 0-back working-memory task blocks, subjects were requested to determine whether the current stimulus matches the target that was presented in the beginning of each block (cue). A 2.5 s cue indicates the task type (and target for the 0-back task) at the beginning of each block. Each of the two sessions contains 8 task blocks (10 trials of 2.5 s each, for 25 s) and 4 fixation (“rest”) blocks (15 s). On each trial, the stimulus is presented for 2 s, followed by a 0.5-s inter-trial-interval (ITI).

### fMRI data acquisition

For each individual, 405 frames were acquired in each session using multiband, gradient-echo planar imaging with the following parameters: TR = 720 ms; TE = 33.1 ms; multiband factor = 8; flip angle = 52°; field of view = 280 × 180 mm; matrix = 140 × 90; and voxel dimensions = 2 mm isotropic.

### fMRI preprocessing

Raw fMRI data for both sessions were obtained from the HCP and underwent standard preprocessing steps, including realignment, slice-time correction, coregistration, normalization, and spatial smoothing with a Gaussian kernel of 6-mm full width at half maximum (FWHM) using SPM12^89^.

### Brain atlas

We leveraged two different atlases, DiFuMo atlas ^60^, and Brainnetome atlas ^102^, to test scalability and robustness of MDSI-AVI in estimating brain-wide causal dynamics. Both atlases provide fine-grained brain-wide parcellations of both cortical and subcortical areas. The DiFuMo atlas ^60^ provides multi-scale functional networks from 64 to 1024 dimensions, allowing us to test the scalability of MDSI-AVI in estimating whole-brain causal interactions, particularly across various number of ROIs. In contrast, the Brainnetome Atlas ^102^ provides superior anatomical and functional interpretability compared to most other atlases. This feature allowed us to investigate brain-wide dynamic circuit mechanisms in human working memory.

### Time series extraction

The original time series were extracted from the pre-processed fMRI data for each ROI, resulting in a matrix with a dimension of T × N, where T is the number of time points and N is the number of voxels in the ROI. Singular value decomposition was applied on the ROI time series matrix, and the resultant first eigenvariate corresponding to the first principal component is obtained to represent the signals of interest within the ROI. The output was a T × 1 vector. We used the first eigenvariate instead of mean signal within the ROI to reduce noise in potentially heterogeneous ROIs. A multiple linear regression approach with 6 realignment parameters (3 translations and 3 rotations) was applied to the time series to reduce head-motion-related artifacts and the data were high-pass filtered (>0.008 Hz).

### Stability analysis

To evaluate robustness of MDSI findings, we conducted stability analysis using a bootstrapping procedure. Stability analysis for multivariate dynamic causal interaction patterns was performed using the following steps:

1. Randomly select a subset of samples from 728 participants without replacement.
2. Apply t-test on each connection per condition or paired t-test on each connection between task conditions.
3. Threshold dynamic causal interaction matrix at p = 0.01 (FDR correction).
4. Repeat steps 1–3 500 times.
5. Compute averaged thresholded dynamic causal interaction matrices.
6. Compute the correlation between the averaged thresholded dynamic causal interaction matrix and the matrix from the original full sample.
7. Change subsample size from 20 to 600 and repeat steps 1–6. The subsample sizes include 20, 30, 40, 50, 60, 70, 80, 90, 100, 150, 200, 250, 300, 400, and 600.
8. Generate stability graph by plotting correlation coefficients from step 6 over subsample size.

### Comparisons with Regression DCM (rDCM)

rDCM is a scalable variant of DCM that leverages the linearity of the model in the frequency domain to accelerate the inference of the directed connectivity matrix. rDCM assumes that the latent activation *x* ∈ *R^M^*^×*T*^ follow the dynamics 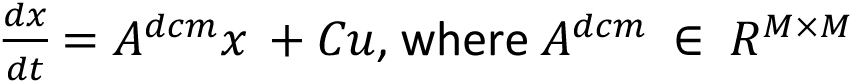 is the connectivity matrix, *C* ∈ *R^M^*^×*M*^ the driving inputs and *u* ∈ *R^M^*^×*T*^ the experimental manipulations impacting each region. *M* and *T* refer to the number of regions and the number of time-steps in **x**, respectively. For experiments using the HCP n-back working memory dataset, the BOLD data was split with respect to conditions. More precisely, considering *Y* ∈ *R^M^*^×*T*^ the BOLD signal for one subject, we derived three *Y_c_* ∈ *R^M^*^×*T*^*^c^* matrices, with *T_c_* the total number of time-steps during which the observed condition was *c*. We constrained that Σ*c*=1*^C^ T_c_* = *T*. Each matrix *Y_c_* was used for the inference of a connectivity matrix *A^dcm^*, defining *u* as an empty vector. rDCM implementation is available in MATLAB as part of the open-source TAPAS toolbox ^103^.

### Comparisons with MDSI-AVI with canonical HRF

To analyze the effect of marginalizing the uncertainty of the HRF, we reported the results of MDSI-AVI assuming the canonical HRF for all regions, or equivalently, fixing *α_m_* = 0 for all *m* ∈ {1, *M*}.

### Test-retest reliability analysis at the group-level and subject-level

We first examined the consistency of whole-brain causal interaction patterns across sessions at the group level. We computed the mean causal interaction patterns for each task (rest, 0-back, and 2-back) across all participants and then assessed the Pearson correlation between group-level estimates from Session 1 (“test”) and Session 2 (“retest”). Next, we investigated the consistency of whole-brain causal interaction patterns across sessions at the individual subject level. Specifically, we tested whether the correlation of whole-brain causal interaction patterns across sessions is higher within subjects than between subjects. For each participant, we computed the Pearson correlation between their connectivity matrix in Session 1 (“test”) and the matrices of all participants in Session 2 (“retest”). Given subject *s*^1^ in Session 1, we identified a subject *s*j^2^ in Session 2 with the highest correlation to *si*^1^. A score *k_i_* is assigned a value of one if *i* = *j* (i.e., the highest correlation is with the same individual), and zero otherwise. Identification accuracy is computed as the average of these scores across all subjects,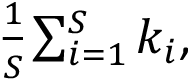 where *S* is the total number of subjects.

### Task classification

To examine the consistency of directed connectivity across conditions, we performed task classification using a logistic regression classifier with L2 regularization to distinguish between three task states (rest, 0-back, and 2-back) based on subject-level causal matrices (*A_c_*). For each subject, causal matrices derived from MDSI-AVI were obtained for all tasks and reshaped into 1D feature vectors. These vectors were standardized and used to train the classifier. To ensure robust performance estimates while accounting for repeated measures within subjects, we applied a 20-fold GroupKFold cross-validation scheme, stratified by subject ID. Model performance was evaluated using confusion matrices, and the final classification score was computed by averaging the true positive rates across the three task conditions. Task classification performance based on MDSI-AVI outputs was compared to those obtained using *Pearson* correlation and rDCM-derived matrices.

### Region-to-Functional Network Mapping

To enable system-level analyses of brain-wide causal interactions, we assigned each brain region in both the DiFuMo and Brainnetome atlases to one of seven canonical functional networks. Specifically, we adopted Yeo et al.’s 7-network parcellation framework, which includes the visual (VIS), somatomotor (SOM), dorsal attention (DAN), ventral attention/salience (SN), limbic (LIM), frontoparietal control (FPN), and default mode (DMN) networks. In addition to these seven cortical networks, we introduced a subcortical (SUBC) network category to account for subcortical regions, which are not explicitly labeled in Yeo’s original framework. We note that not all network labels are present at all parcellation scales. Notably, in DiFuMo atlas, regions assigned to the limbic system (LIM) are only present in the 256-region version of the atlas, but not in the 64- or 128-region versions used in our main analyses, including Figure 6. As a result, the LIM network was excluded from those analyses.

### Direct and indirect causal pathway analysis

To examine how the right dorsal anterior insula (dAI) influences network dynamics during working memory, we analyzed both direct and indirect causal pathways originating from the right dAI to other brain networks. A direct and indirect pathway from a target region A (in our case, right dAI) to regions B and C is defined as AB(direct)C(indirect) when the causal weight from A to C is not significantly different from zero (FDR-corrected, *p*<0.05), while the causal weights from A to B and from B to C are significant. By testing all possible combinations of regions B and C, we estimated both direct and indirect causal pathways originating from region A. We then assigned each region comprising B and C to one of eight intrinsic functional systems ^61^ and computed the average causal weight from region A. Through this process, we estimated how information potentially propagates from the right dorsal anterior insula (dAI) through multiple pathways in the brain, providing insights into the neural processes underlying working memory.

### Brain-behavior analysis

We applied canonical correlation analysis (CCA) ^104^ to explore the relation between dynamic causal interactions and working-memory performance. CCA is a statistical method for examining the relationships between two multivariate sets of variables, and has been shown to be a powerful tool for investigating brain-behavior relationships. CCA finds the optimal linear combination of subjects’ multivariate behavioral measures that maximize the relation between behavioral and brain measures. Specifically, brain features included dynamic system-level causal interaction weights of connections in 0-back or 2-back conditions, and behavioral features included accuracy and reaction time in 0-back or 2-back conditions. The significance of the canonical relationship was tested using permutation testing. CCA was rerun after each permutation, and the canonical correlation was recalculated. These procedures were repeated 1,000 times to build a null distribution of canonical correlations for comparison with the original canonical correlation. The p-values of each CCA dimension were corrected for multiple testing across the estimated CCA modes (FDR-corrected, p < 0.05).

## DATA AVAILABILITY

All original code used for the analysis and all data utilized in this study will be deposited online and made available without restrictions.

## ACKNOWLEDGMENTS

This research was supported by grants from the National Institute of Health (NS086085, MH126518 to V.M.), the Alzheimer’s Association (AARFD-21-848178 to B.L. and AARGD-NTF-21-850781 to W.C.), and the Korea Institute of Science and Technology (G04250061 to B.L).

## AUTHOR CONTRIBUTIONS

Conceptualization: BL, LR, SR, DW, VM; Methodology: BL, LR, LA, SR, DM, VM; Data acquisition: BL, WC; Investigation: BL, LR, LD, LJ; Writing-original draft: BL, VM; Writing-review&editing: BL, LR, LD, LJ, LA, SR, NB, PM, WC, DW, VM.

## COMPETING INTERESTS

The authors declare no competing interests.

## Notes

### Competing Interest Statement

The authors have declared no competing interest.

